# Naphthalimide-based, Single-Chromophore, Ratiometric Fluorescent Sensor for Tracking Intracellular pH

**DOI:** 10.1101/2024.05.14.594096

**Authors:** Sujit Kumar Das, Smitaroopa Kahali, Sabnam Kar, Nandita Madhavan, Ankona Datta

**Author notes:** Electronic supplementary information (ESI) available: Computational methods, additional absorbance and fluorescence studies, experimental details of cell studies are in Supporting Information. These authors contributed equally (Sujit Kumar Das, Smitaroopa Kahali).

## Abstract

We report a novel, reversible, cell-permeable, pH-sensor, TRapH. TRapH afforded a pH-sensitive ratiometric emission response in the pH range ∼3-6, enabling imaging and quantification of pH in living cells. The biological-applicability of TRapH was illustrated via live-tracking of intracellular pH dynamics in living mammalian cells induced by a synthetic H^+^-transporter.

---

Intracellular pH plays a central role in maintaining cellular homeostasis.^1^ In healthy mammalian cells the cytoplasmic pH lies between 7.0-7.4.^1a, 2^ Intracellular organelles have different pH ranges tailored to their functions.^1a^ For example, the pH range is 4.5-5.0 in lysosomes to allow waste degradation,^3^ and 7.8-8.2 within mitochondria to facilitate ATP synthesis and enable mitochondrial enzyme activity.^1a, 4^ Deviations in cellular pH are associated with pathophysiological conditions including cancers^5^ and neurological disorders.^6^ Therefore, intracellular pH is a critical parameter that determines the physiological condition of a cell. In this context, pH sensitive fluorescent probes are valuable non-invasive tools for monitoring intracellular pH dynamics and for relating this vital parameter to pathophysiological states.^7^

Key requirements for non-invasive, instantaneous pH detection in living cells are rapid-response fluorescence probes that can report on pH changes within the physiologically relevant pH range of 4-8. An ideal pH probe for cellular pH detection should afford visible excitation and emission, and a ratiometric response for pH quantification. Herein, we have developed a photostable, ratiometric pH probe, TIFR Ratiometric pH sensor (**TRapH)**, based on a naphthalimide dye.

The reason for choice of naphthalimide was its high photo-stability, tuneable optical properties, and ease of substitution toward introduction of responsive functional groups.^8^ Unlike the few previous reports on naphthalimide-based pH sensors which use piperazine as the pH sensitive unit,^8b, c^ we decided to use alkyl-aniline as our pH sensitive unit. This is because, we noted that the first pKa of alkyl-piperazine is ∼ 9.7^9^ while that of alkyl-aniline is ∼5.5-6.6.^10^ Hence, based on the expectation that a pH probe would respond within ± 1 pH range of the pKa value of its pH sensitive unit, we reckoned that alkyl-anilines would be an appropriate choice. Importantly, alkyl-anilines would be able to cover almost the entire physiologically relevant pH range without the requirement of either an additional pH sensitive unit or an additional pH responsive dye as has been the case for previously reported naphthalimide-based pH sensors.^8b, d^ We therefore zeroed in on the blue-print for **TRapH** wherein a diethylaniline moiety was ‘clicked’ to a naphthalimide unit (Fig 1a).

**Fig. 1.**
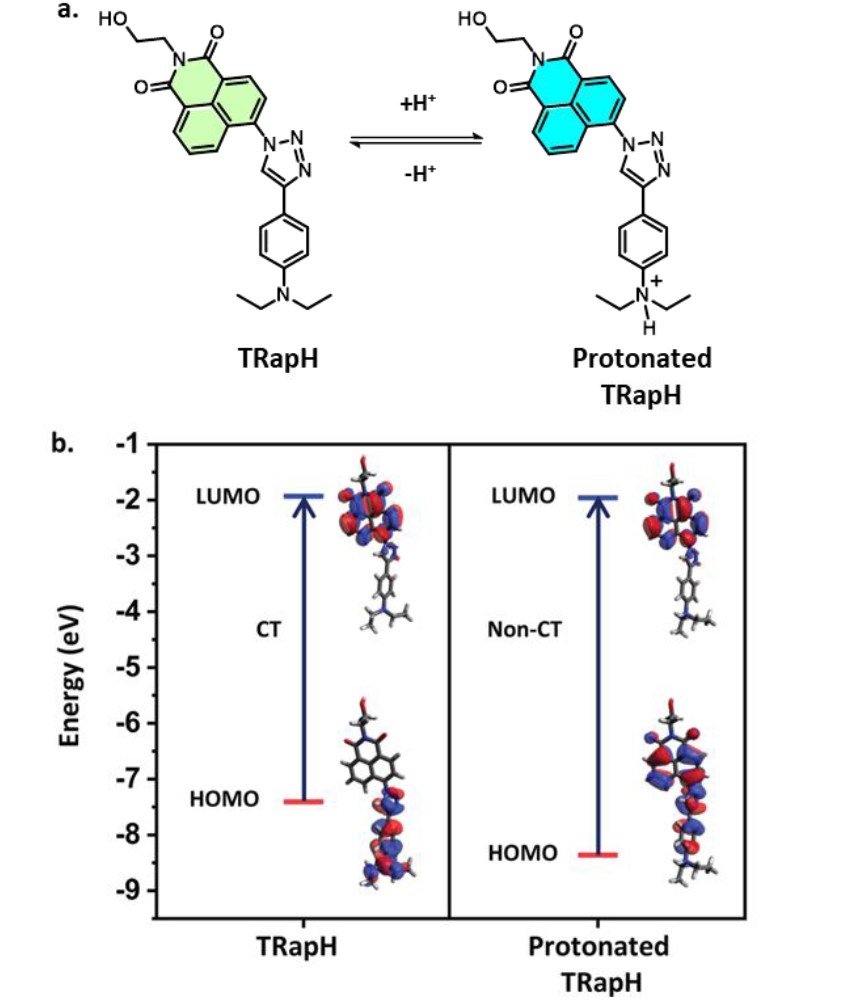
(a) Schematic showing mechanism of action of **TRapH**. (b) Plots depicting the energy levels of optimised ground state geometries along with representative transitions with highest oscillator strengths in **TRapH** and protonated **TRapH**.

Before proceeding with the synthesis and evaluation of the designed probe, we decided to perform electronic structure calculations to predict the photophysical properties of **TRapH**. A computational framework to predict the photophysical properties of the emerging class of naphthalimide-based sensors would not only help guide our experimental studies but also enhance understanding of this dye-class and enable improved design. We computationally investigated the effect of protonation of the diethylaniline moiety on the electronic transitions in **TRapH** (Fig 1b, S1, S2, Table S1 and S2). The goal was to predict whether and how **TRapH** might act as a pH sensor. Density functional (DFT) and time-dependent density functional (TD-DFT) calculations were performed on **TRapH** and protonated **TRapH** both in vacuum and in a suitable dielectric medium, here water (Fig 1b, S1, S2, Table S1 and S2). **TRapH** showed a distinct intramolecular charge transfer (ICT) transition between diethylaniline and the naphthalimide dye core (Fig 1b, S1 and Table S1) and multiple non-CT transitions (Fig S1 and Table S1). We noted that in protonated **TRapH** the ICT transition was completely absent (Fig 1b and Table S2). The major transitions involved orbitals delocalized over the dye, triazole, and phenyl ring (Fig 1b, S2 and Table S2). Based on the computational studies, we hypothesized that the absence of the ICT transition in protonated **TRapH** might translate into altered emission with variation in pH.

We next proceeded to validate the computational results experimentally. **TRapH** was synthesised and characterized. pH dependent absorption spectra of **TRapH** were recorded in aqueous buffers with pH range 2-8 (Fig 2a). **TRapH** showed two absorption peaks at neutral to basic pH (pH 7.4-8), one at 298 nm (18420 ± 538 M^-1^ cm^-1^) with a shoulder at 334 nm, and the other at 438 nm (2060 ± 22 M^-1^ cm^-1^) (Fig 2a and S3). On lowering the pH, a new peak emerged at 347 nm (13500 ± 219 M^-1^ cm^-1^) (Fig 2a and S3) and the absorbance at 438 nm decreased significantly (Fig 2a). The absorption band centred at 438 nm was broad and in the visible region which indicated that this absorption feature could correspond to an ICT transition. As per the computational studies, the ICT transition would be prominent in **TRapH** and absent in protonated **TRapH** (Fig 1a). The absorption spectra also showed the same trend.

**Fig. 2.**
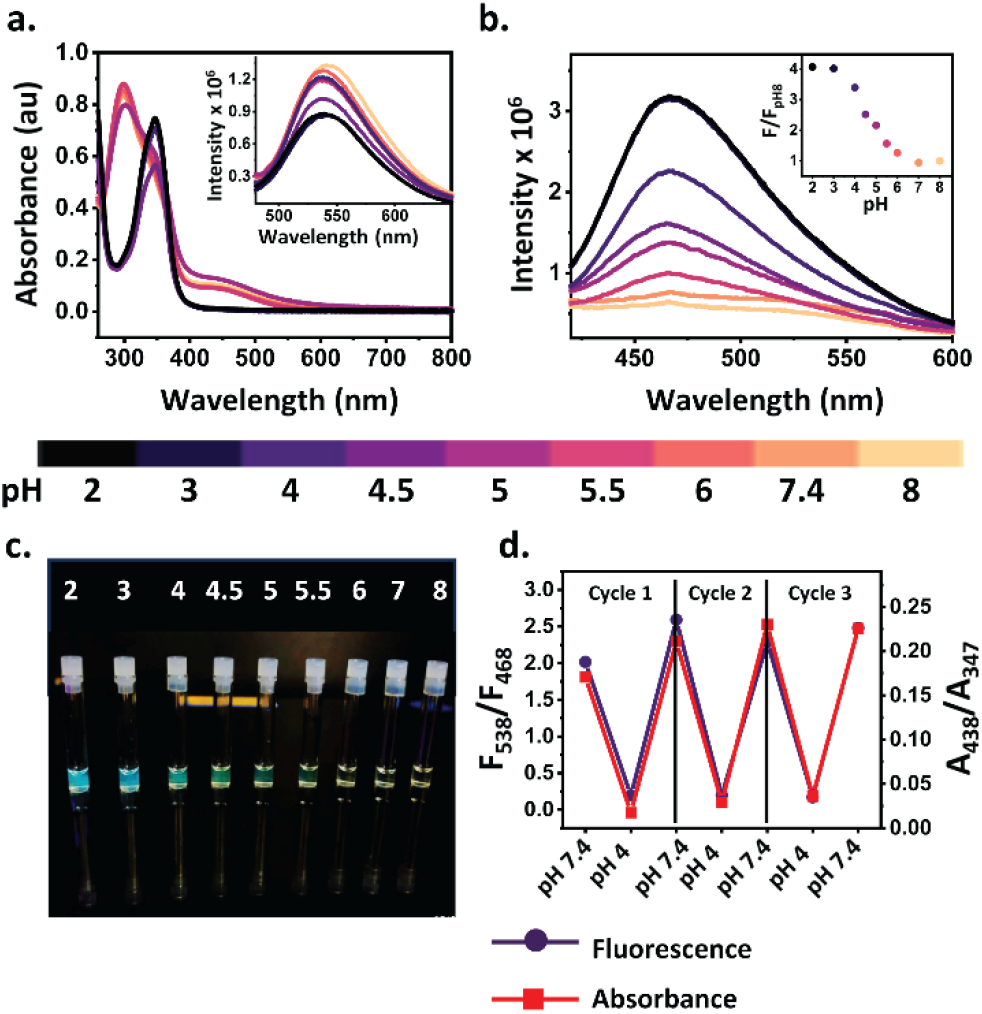
(a) pH-dependent absorption spectra of **TRapH** (50 μM). Inset: pH-dependent fluorescence response of **TRapH** (50 μM); λ_ex_ = 450 nm. (b) pH-dependent fluorescence response of **TRapH** (50 μM); λ_ex_ = 347 nm. Inset: Variation of ratio of fluorescence (F) over fluorescence at pH 8 (F_pH8_) with pH; λ_ex_ = 347 nm (c) Image depicting vials containing **TRapH** (50 μM) in buffers of different pH under a long-wavelength UV lamp (λ_ex_ = 365 nm). (d) Reversible changes in absorption (red) and fluorescence (purple) of **TRapH** with alternating pH, acidic (pH 4) to neutral (pH 7.4).

To confirm whether the 438 nm absorption band was an ICT transition, absorption spectra for **TRapH** were recorded in solvents with different polarities (Fig S4a). The peak showed a solvent-dependent shift (Fig S4a). We next recorded fluorescence spectra in different solvents. Upon excitation at 450 nm, an 88 nm stokes-shifted emission peak was observed with a maxima at 538 nm in aqueous buffer (Fig 2a inset) which also showed a distinct solvent-dependent shift for the peak maxima (Fig S4b). The solvent-dependent shift in both absorption and emission spectra indicated that the 438 nm absorption peak indeed corresponded to an ICT transition. Importantly, the absorbance at 438 nm decreased with decreasing pH while the absorbance at 347 nm increased with decreasing pH (Fig 1a). As a result, the ratio of absorbances at 438 nm to 347 nm decreased with decreasing pH (Fig S5) indicating that **TRaPH** could be developed as a ratiometric pH sensor.

To investigate whether the observed absorption changes would lead to a pH-dependent fluorescence response, emission spectra for **TRapH** were recorded in aqueous buffers of pH range 2-8 (Fig 2b). 450 nm and 347 nm were used as excitation wavelengths. The intensity of the emission peak at 538 nm (λ_ex_ 450 nm) corresponding to the CT band excitation showed 1.5 times decrease upon lowering the pH (Fig 2a inset). On the other hand, upon excitation at 347 nm, **TRapH** exhibited low fluorescence emission with a maxima at 468 nm at neutral to slightly basic pH (7.4-8) which increased 4-fold on lowering the pH (Fig 2b). The response saturated at pH 3 (Fig 2b inset). The absorbance versus concentration plots could be fitted to straight-lines in the concentration range used for all our studies (Fig S3). We therefore, eliminated any possibility of aggregation induced emission and concluded that the pH-dependent fluorescence response was due to changes in absorption features. Since, the two fluorescence emission bands afforded opposite response to pH, we concluded that **TRapH** could act as a ratiometric fluorescent pH sensor.

We next determined the pKa of **TRapH** by two methods: 1. from pH dependent absorption data by using Henderson-Hasselbach equation (Fig S6a); 2. by determining the mid-point of the sigmoidal pH-dependent fluorescence emission data. (Fig S6b). The pKa for **TRapH**, calculated from absorption data, was 4.6 ± 0.2 (Fig S6a) and from fluorescence data was 4.6 (Fig S6b). The pKa value for **TRapH** was lower than that reported for diethylaniline possibly due to conjugation of aniline with the triazole unit. A pKa of 4.6 indicated that **TRapH** would be able to sense pH changes within the pH range of ∼3-6. An image of vials containing **TRapH** excited under a long UV lamp (365 nm) also validated that the range within which **TRapH** would effectively report pH, was between 3-6 (Fig 2c).

Reversibility of absorption and fluorescence response allows detection of dynamic changes in pH. To determine if **TRapH** was a reversible pH probe, absorbance and fluorescence spectra were measured by dissolving the sensor in aqueous buffer and alternately adding HCl and NaOH to cycle the pH between pH 4 and 7.4, respectively (Fig 2d). Spectra were recorded after each addition. Both, absorbance and fluorescence responses of **TRapH** were reversible for three cycles of pH variation (Fig 2c), indicating that **TRapH** was a reversible, ratiometric, fluorescent pH sensor.

To further characterize pH dependent structural and electron-distribution changes in **TRapH**, ^1^H NMR spectra were recorded at pH 7.5 and 3.5 (Fig 3 and S7). Protonation of the diethylaniline amine moiety should lead to a downfield shift of protons at positions directly conjugated to the aniline-*N* lone pair. We indeed observed a 0.74 ppm shift for the *ortho-* proton of diethylaniline and a 1.25 ppm shift for the triazole proton. Most of the naphthalimide protons also showed a downfield shift. This might be attributed to a lower electron-density at the dye core due to the electron-withdrawing inductive effect of the positively-charged anilinium moiety. The ^1^H NMR spectra confirmed that the pH sensitive response of **TRapH** was due to diethylaniline protonation. Finally, we tested the applicability of **TRapH** toward intra-cellular pH detection in living mammalian cells. Since probe excitation in the UV, at ∼350 nm, would not be conducive for live cell imaging, 405 nm excitation was selected for the cell imaging studies. Both **TRapH** and protonated **TRapH** could be excited at 405 nm (Fig S8) since both species showed some absorption at this wavelength. Before proceeding with the in-cell studies, pH dependent fluorescence spectra for **TRapH** were recorded in vitro, via excitation at 405 nm (Fig S9). Both the 468 nm and 538 nm emission bands were observed (Fig S9a). The ratio of integrated fluorescence response in blue (430-475 nm) and green (510-600 nm) regions afforded the same trend with pH as observed previously for the fluorescence spectra obtained with excitation at 347 nm (Fig S9b). Based on this result, living HeLa cells were incubated with **TRapH** at pH 7.4. The sensor permeated living cells within 5 min direct incubation (Fig S10 and S11). In accordance with an expected intracellular pH of ∼7.0-7.4, we observed significantly low fluorescence from the blue channel and high fluorescence from the green channel upon excitation at 405 nm in a confocal microscopy set-up (Fig S10 and S11).

**Fig. 3.**
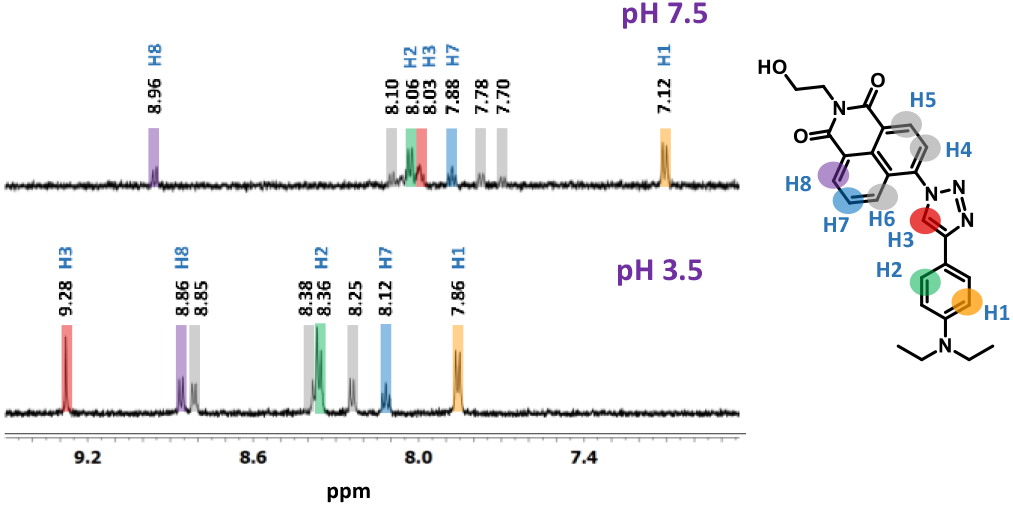
Aromatic region of ^1^H NMR spectra of **TRapH** recorded at two different pH, pH 7.5 (top panel) and pH 3.5 (bottom panel).

Since **TRapH**, showed a ratiometric response between a pH range of 3-6, we next decided to apply a previously established peptide-based synthetic H^+^ ion transporter, **Amphiphilic Peptide 1a** (Fig S12),^11^ to HeLa cells to reduce intracellular pH, and record temporal pH changes live (Fig 4). **Amphiphilic Peptide 1a**, has been reported to reduce pH inside vesicles via transport of H^+^ ions.^11^ A time-lapse imaging experiment in the absence of the peptide at pH 7.4 showed no change in emission proving the photostability of **TRapH** (Fig S13). For imaging temporal pH changes, HeLa cells were first incubated with **TRapH** (5 min), washed, and then incubated with **Amphiphilic Peptide 1a** for 45 min at pH 7.4. An image was recorded on a confocal microscopy set-up at the end of 45 min (Fig 4a, top panel). Following that, the media was removed carefully and media at pH 4.5 was added to the cells (Fig 4c). A gradual increase in the blue channel intensity and a concomitant decrease in the green channel intensity were observed when a time-lapse image was recorded for 20 min at intervals of 1 min (Fig 4a bottom panel, 4b). The average ratio of the mean fluorescence intensities in the blue and the green channel per cell, saturated at 0.5 (Fig 4b). From the in vitro calibration curve (Fig S9b), we concluded that the intracellular pH had stabilized at 4.5. These results indicated that **TRapH** was a photostable, visible-excitable, ratiometric, cell-permeable, fluorescent sensor that could track and quantify pH dynamics in living cells.

**Fig. 4.**
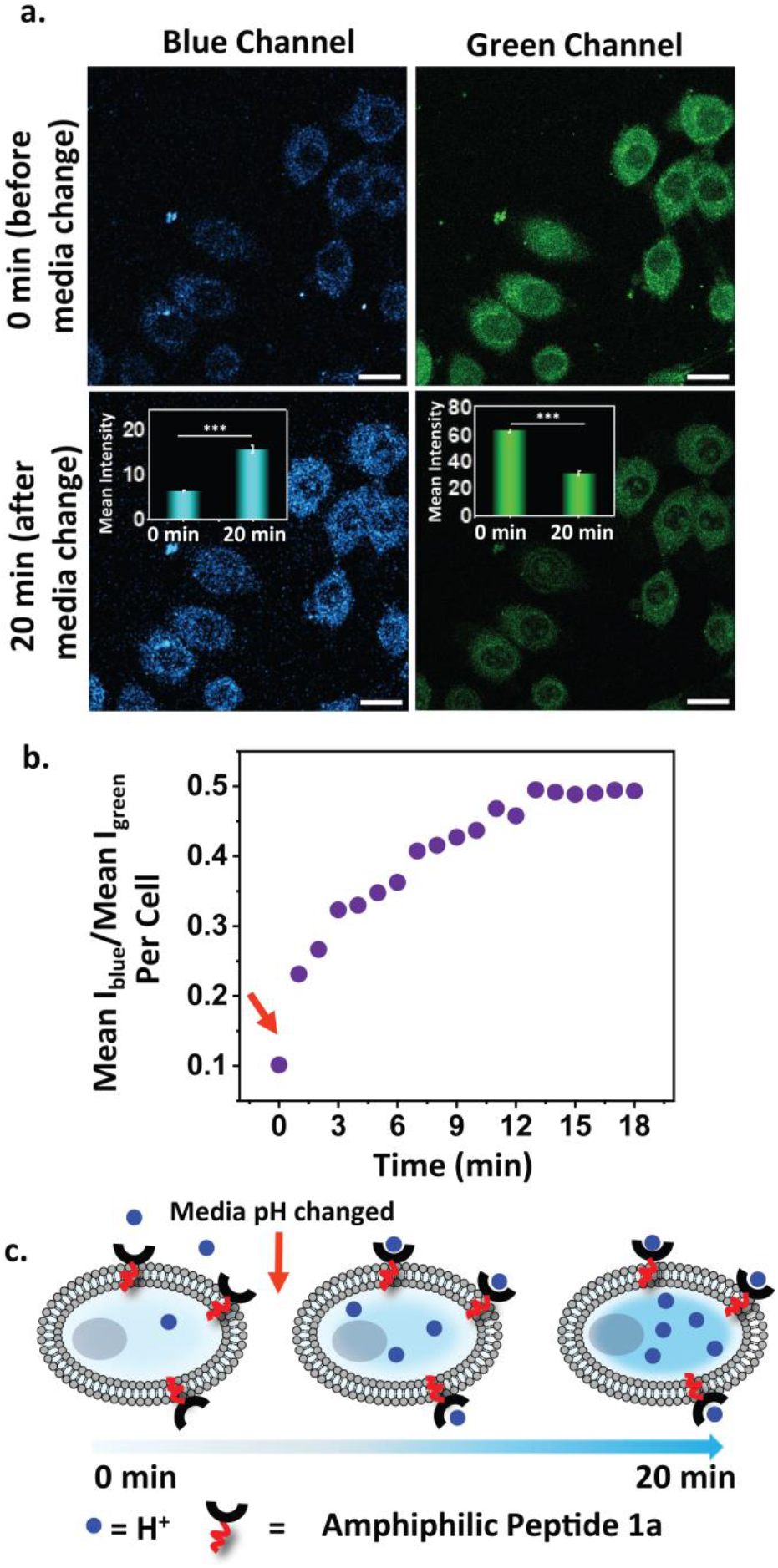
(a) Representative confocal single z plane images of HeLa cells in blue (430-475 nm) and green channels (510-600 nm). λ_ex_ = 405 nm, scale bar = 20 μm. HeLa cells incubated with **TRapH** (10 μM, pH 7.4) followed by **Amphiphilic Peptide 1a** (10 μM, pH 7.4). After 45 min, an image was recorded (top panel). Media was replaced with media at pH 4.5, and time-lapse images were recorded. Image at 20 min shown in bottom panel. Inset: Bar plots representing mean intensities obtained from intensity analysis of images shown in top and bottom panels. Data are presented as SEM, where *n* = 10 for each set. \**p* < 0.05, \*\**p* < 0.01 and \*\*\**p* < 0.001. (b) Tracking pH dynamics. Plot of ratio of mean intensities in blue and green channels tracked at intervals of 1 min for 20 min, starting from the time of change to pH 4.5 media (indicated by arrow). (c) Schematic depicting the experimental flow for tracking intracellular pH dynamics.

Thus far, most of the few reported ratiometric naphthalimide-based pH probes have employed two dyes for achieving a ratiometric response.^8b, d^ **TRapH** is a single chromophore-probe that affords pH-sensitive ratiometric response within most of the physiologically-relevant pH range. Importantly, a key pH-dependent photophysical property of **TRapH** could be predicted via computational studies before its synthesis which highlights the advantage of utilizing computations in molecular probe design.

## Acknowledgements

A. D. acknowledges support from DAE, Govt. of India, Project No. RTI4003. Sabnam Kar acknowledges DST/WOS-A/CS-133/2021 and NM acknowledges SERB (CRG/2022/007764). The authors acknowledge Prof. Vamsee. K. Voora (TIFR) for discussions on computations; National NMR facility, and DCS Cell Culture Facility.

## Notes and references

1. (a)J. R. Casey, S. Grinstein and J. Orlowski, Nat. Rev. Mol. Cell Biol., 2010, 11, 50–61; (b) M. Tantama, Y. P. Hung and G. Yellen, J. Am. Chem. Soc., 2011, 133, 10034–10037.

2. I. H. Madshus, Biochem. J., 1988, 250, 1–8.

3. R. Chen, M. Jäättelä and B. Liu, Cancers, 2020, 12.

4. M. F. C. Abad, G. Di Benedetto, P. J. Magalhães, L. Filippin and T. Pozzan, J. Biol. Chem., 2004, 279, 11521–11529.

5. S. Lee and A. Shanti, Int. J. Mol. Sci., 2021, 22.

6. T. I. Lam, A. M. Brennan-Minnella, S. J. Won, Y. Shen, C. Hefner, Y. Shi, D. Sun and R. A. Swanson, Proc. Natl. Acad. Sci. U.S.A., 2013, 110, E4362–E4368.

7. (a)K.-K. Yu, J.-T. Hou, K. Li, Q. Yao, J. Yang, M.-Y. Wu, Y.-M. Xie and X.-Q. Yu, Sci. Rep., 2015, 5, 15540; (b) F. Le Guern, V. Mussard, A. Gaucher, M. Rottman and D. Prim, Int. J. Mol. Sci., 2020, 21; (c) A. Maity, U. Ghosh, D. Giri, D. Mukherjee, T. K. Maiti and S. K. Patra, Dalton Trans., 2019, 48, 2108–2117; (d) Y. Wu, C. Ge, Y. Zhang, Y. Wang and D. Zhang, Front. Chem., 2023, 11; (e) S. Das, A. Kapadia, S. Pal and A. Datta, ACS Sens., 2021, 6, 2252–2260; (f) M. Martineau, A. Somasundaram, J. B. Grimm, T. D. Gruber, D. Choquet, J. W. Taraska, L. D. Lavis and D. Perrais, Nat. Commun, 2017, 8, 1412; (g) J. Qi, D. Liu, X. Liu, S. Guan, F. Shi, H. Chang, H. He and G. Yang, Anal. Chem., 2015, 87, 5897–5904.

8. (a)N. I. Georgiev and V. B. Bojinov, J. Lumin., 2012, 132, 2235–2241; (b) X. Zhou, F. Su, H. Lu, P. Senechal-Willis, Y. Tian, R. H. Johnson and D. R. Meldrum, Biomaterials, 2012, 33, 171–180; (c) K. Pršir, M. Matić, M. Grbić, G. J. Mohr, S. Krištafor and I. M. Steinberg, Molecules, 2023, 28; (d) P. Srivastava, P. Srivastava and A. K. Patra, New J Chem, 2018, 42, 9543–9549; (e) S. Zheng, P. L. M. Lynch, T. E. Rice, T. S. Moody, H. Q. N. Gunaratne and A. P. de Silva, Photochem. Photobiol. Sci., 2012, 11, 1675–1681.

9. F. Khalili, A. Henni and A. L. L. East, J. Chem. & Eng. Data., 2009, 54, 2914–2917.

10. G. A. Gohar and M. M. Habeeb, Spectroscopy, 2000, 14, 831496.

11. S. Kar and N. Madhavan, Chem. Eur. J., 2023, 29, e202301020.

