## supporting information for "Naphthalimide-based, Single-Chromophore, Ratiometric Fluorescent Sensor for Tracking Intracellular pH"

‡ Equal contribution

Corresponding author: Ankona Datta

**General Procedures and Methods**

All chemicals used were of analytical grade, obtained from commercial sources and used without further purification unless otherwise mentioned. Chemicals were purchased from either Sigma Aldrich®, or SD Fine-Chem Ltd., or TCI India Research Chemicals, or Arrobiochem Private limited and used without further purification unless otherwise noted. All cell culture reagents were purchased from either Sigma Aldrich® or Gibco®. Analytical grade solvents were used for synthesis and separation, and used without distillation.

HeLa cells were purchased from American Type Culture Collection (ATCC®).

UV-Visible spectrophotometric experiments were performed on a Thermo Scientific Multiskan Go spectrophotometer in a quartz cuvette having a path length of 1 cm with 10 mm x 4 mm (Hellma® Analytics) inner dimensions.

Liquid chromatography mass spectrometry (LCMS) analyses were carried out on a Shimadzu LCMS 2020 with an electrospray ionization (ESI) probe (positive and negative ion modes). High-resolution mass spectrometry (HRMS) analysis was carried out on Thermo Scientific™ Q Exactive™ hybrid quadrupole-Orbitrap™ mass spectrometer.

**Geometry optimization of molecules and visualization of electron densities on molecular orbitals (MOs):**

Initial geometries of **TRapH** and protonated **TRapH** were generated using GaussView 6.0.16 software. DFT calculations were performed on the initially generated geometries of the molecules to obtain the energy optimized ground state geometries of the respective molecules in vacuum and in a suitable dielectric medium (water) using CAM-B3LYP correlation functional and 6-311++G$^{**}$ basis set in Gaussian 16 software. Solvent was modeled using polarizable continuum model (PCM); no explicit solvent molecules were added during the calculation. Molecular orbitals (MOs) of the molecules were visualized using Avogadro 1.1.1 software.

**Identifying potential transitions after photo-excitation:**

Time-dependent density functional theory (TD-DFT) was applied on the energy optimized ground states of **TRapH** and protonated **TRapH** to determine the most probable transitions in the molecules in water. Optical transitions with maximum oscillator strength (f) were considered as most probable transitions both in **TRapH** and protonated **TRapH**. Optical transitions in **TRapH** were categorized into two sets (a) Charge transfer (CT) transitions (transitions where electron density on the ligand part i.e. triazole containing diethyl aniline unit gets transferred to the dye part i.e. naphthalimide unit after photo-excitation) (b) Non-

CT transitions where participating orbitals have electron density either on the naphthalimide dye core or delocalized over the whole molecule. Most probable CT and non-CT transitions are noted below along with the respective transition probabilities (TP).

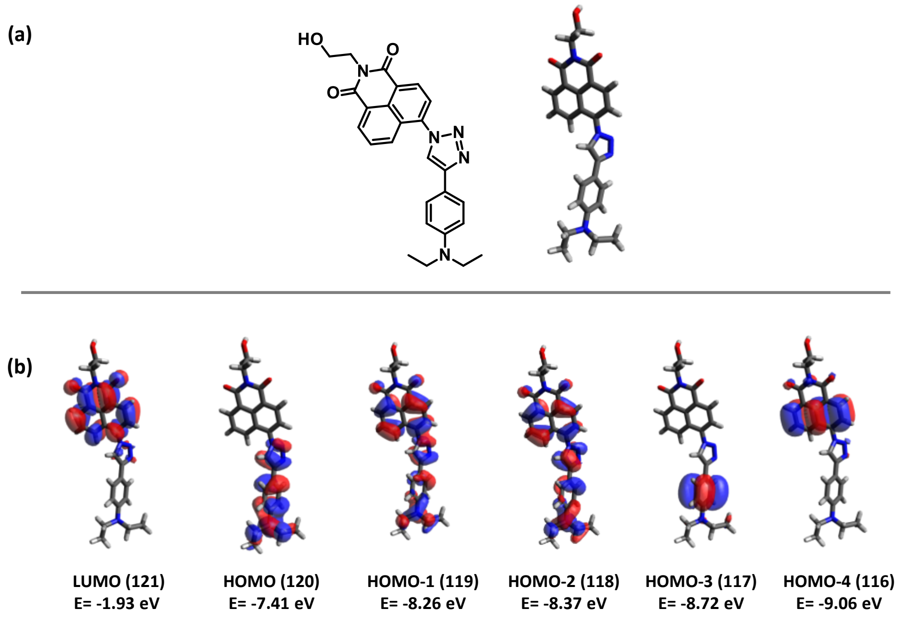


**Figure S1.** (a) Energy optimized ground state geometry of **TRapH** in water**.** (b) Representative molecular orbitals of **TRapH,** relevant to the most-probable transition, in water.

**Table S1.** Optical transitions corresponding to highest oscillator strength in **TRapH**.

| TRapH | | | |
|---|---|---|---|
| **Optical Transitions** | **Transition wavelength (nm)** | **Oscillator strength** | **Transition probability** |
| **HOMO-2 → LUMO (Non CT)** | | | **0.36** |
| **HOMO-1 → LUMO (Non-CT)** | **317** | **0.57** | **0.56** |
| **HOMO → LUMO (CT)** | | | **0.19** |

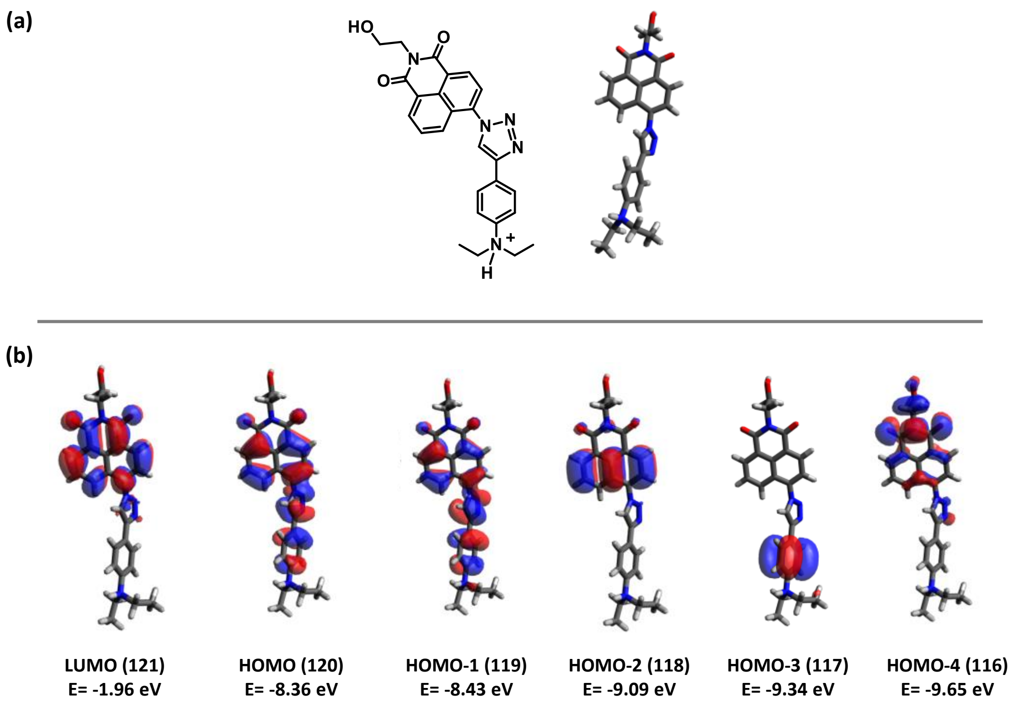


**Figure S2.** (a) Energy optimized ground state geometry of protonated **TRapH** in water. (b) Representative molecular orbitals of protonated **TRapH**, relevant to the most-probable transition, in water.

**Table S2.** Optical transitions corresponding to highest oscillator strength in protonated **TRapH**.

| TRapH | | | |
|---|---|---|---|
| **Optical Transitions** | **Transition wavelength (nm)** | **Oscillator strength** | **Transition probability** |
| **HOMO -1 → LUMO (Non CT)** | **314** | **0.52** | **0.37** |
| **HOMO → LUMO (Non-CT)** | | | **0.58** |

### <u>In vitro absorption measurement of TRapH:</u>

All spectroscopic measurements for **TRapH** were performed in 5% DMSO, aqueous buffer (HEPES, 20 mM) at room temperature. A stock solution of **TRapH** (10 mM) was prepared in DMSO. The stock solution was diluted in aqueous buffer (HEPES, 20 mM) to prepare **TRapH** solution (50 μM) maintaining 5% DMSO in the aqueous buffer. Absorption spectra were recorded on a Thermo Scientific Multiskan GO UV/Vis Microplate Reader Spectrophotometer using quartz cuvettes with 10 mm x 2 mm inner dimensions (Hellma® Analytics). To record the pH dependent response for **TRapH**, 5% DMSO, aqueous buffer (HEPES, 20 mM) of pH values 2, 3, 4, 4.5, 5, 5.5, 6, 7.4, and 8, were prepared either by adding few drops of HCl or NaOH to the buffer solution. **TRapH** (50 μM) solutions in 5% DMSO, aqueous buffer (HEPES, 20 mM) for each pH, were prepared and absorption spectra were recorded. Solvent dependent absorption spectra for **TRapH** (50 μM) were recorded in solvents with different polarities (acetonitrile, methanol, chloroform, dimethylsulfoxide, and aqueous buffer).

**Determination of the extinction coefficients ($\varepsilon$) for TRapH:**

Extinction coefficients for **TRapH** were determined at different wavelengths (Fig. S4) in 5% DMSO-aqueous buffer (HEPES, 20 mM) system at two different pH values, pH 7.4 and pH 4. Absorption spectra for **TRapH** at different concentrations (80 μM, 60 μM, 40 μM, 20 μM) were recorded in 5% DMSO, aqueous buffer (HEPES, 20 mM) at both pH values. Absorbance value at each wavelength were plotted with varying concentrations of **TRapH** and $\varepsilon$ values were obtained from the slope of the linear fit of the absorbance versus concentration plot using the following equation:

$$\varepsilon = A/Cl$$

$\varepsilon$ is the molar extinction coefficient, A is the absorbance at respective wavelength at particular concentration, C is the concentration of the sample used in the experiment and l is the path length for beam travel.

Each experiment was performed in triplicate and mean values have been reported as $\varepsilon$.

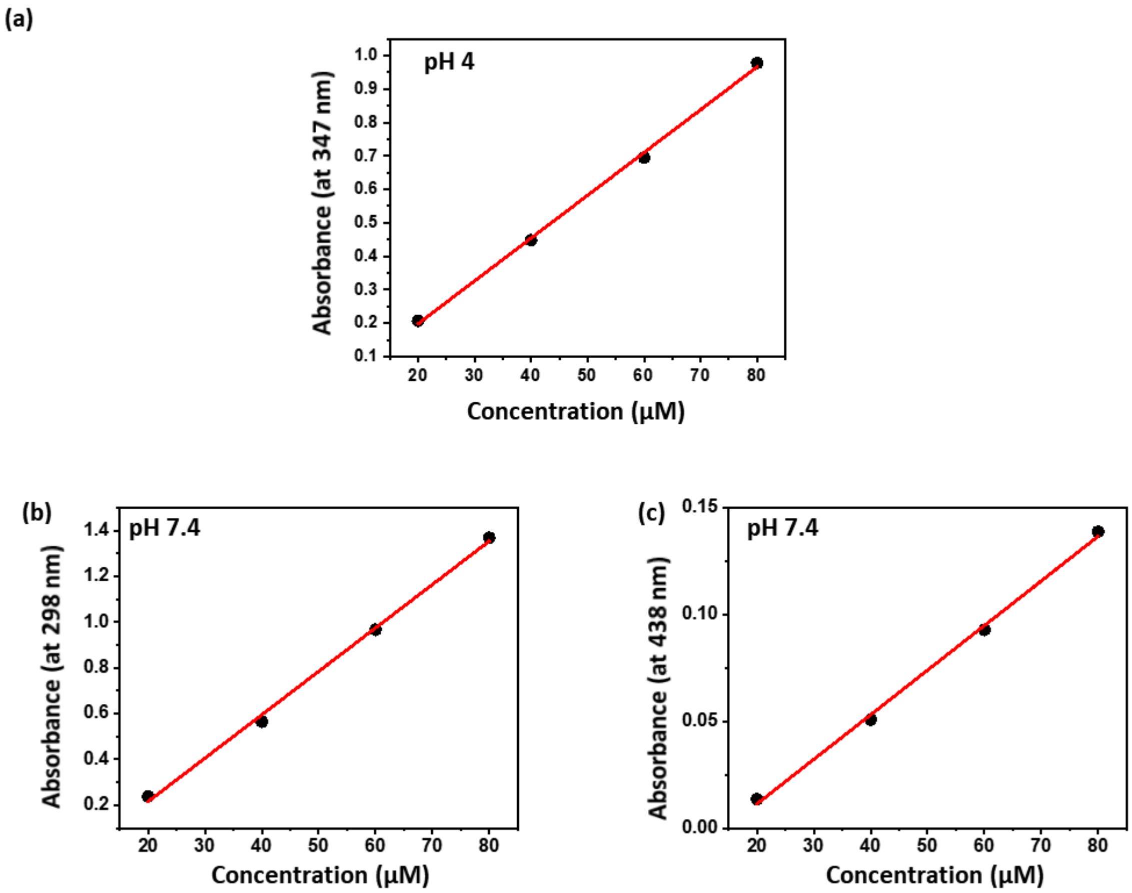


**Figure S3.** Absorbance vs concentration plot: (a) for **TRapH** in 5% DMSO, aqueous buffer (HEPES, 20 mM, pH 4) at 347 nm; (b) for **TRapH** in 5% DMSO, aqueous buffer (HEPES, 20 mM, pH 7.4) at 298 nm; (c) for **TRapH** in 5% DMSO, aqueous buffer (HEPES, 20 mM, pH 7.4) at 438 nm; for calculation of molar extinction coefficient.

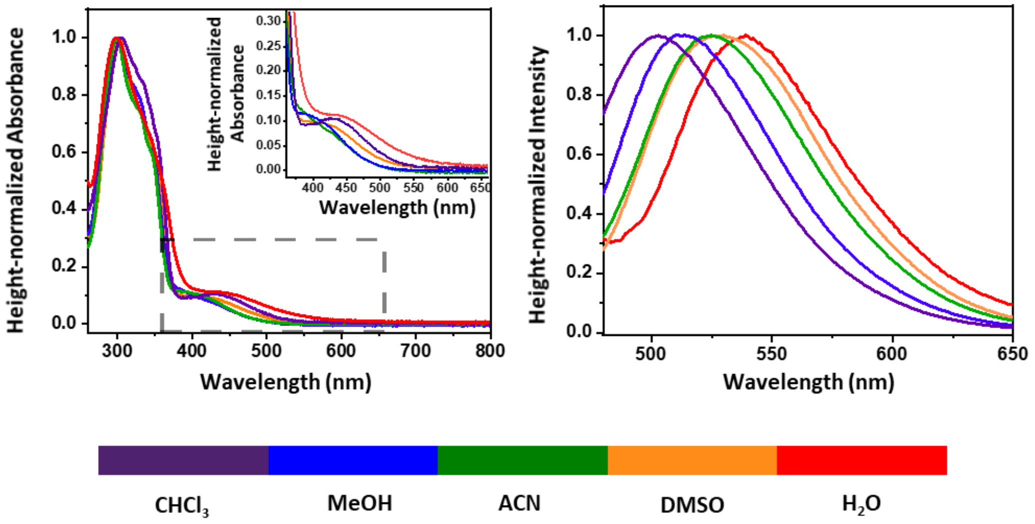


**Figure S4**. (a) Height-normalized absorption spectra of **TRapH** (50 µM) in solvents of different polarities. Inset: Variation in the CT band of **TRapH** with solvent polarity. (b) Height-normalized fluorescence spectra for **TRapH** (50 µM) in solvents of different polarities. $\lambda_{ex}$ = 450 nm, slit width 3 nm x 3 nm.

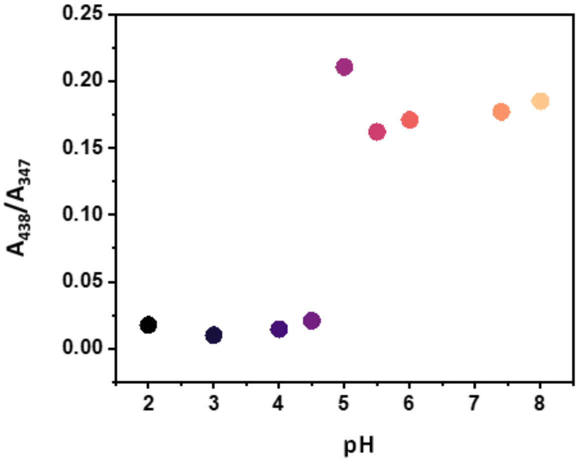


**Figure S5.** Plot depicting the variation of the ratio of absorbance at 438 nm ($A_{438}$) and absorbance at 347 nm ($A_{347}$), with changing pH values.

### <u>In vitro fluorescence measurement of TRapH:</u>

All spectroscopic measurements for **TRapH** were performed in 5% DMSO, aqueous buffer (HEPES, 20 mM) at room temperature. A stock solution of **TRapH** (10 mM) was prepared in DMSO. The stock solution was diluted in aqueous buffer (HEPES, 20 mM) to make **TRapH** solution (50 µM) maintaining 5% DMSO in aqueous buffer (HEPES, 20 mM). Fluorescence spectra were recorded on FluoroLog®-3 (Horiba Jobin Yvon Inc.) spectrofluorometer using quartz cuvettes with 10 mm x 2 mm inner dimensions (Hellma® Analytics). To obtain the pH dependent response for **TRapH**, 5% DMSO, aqueous buffer (HEPES, 20 mM) of pH values 2, 3, 4, 4.5, 5, 5.5, 6, 7.4, and 8 were prepared either by adding few drops of HCl or NaOH. **TRapH** (50 μM) solutions in 5% DMSO, aqueous buffer (HEPES, 20 mM) for each pH were prepared and fluorescence spectra were recorded by exciting at 347 nm and 450 nm, with slit width 4 nm x 4 nm for excitation at 347 nm and 3 nm x 3 nm for excitation at 450 nm. Fluorescence spectra was also recorded by exciting at 405 nm with a slit width of 3 nm x 3 nm. For solvent-dependent emission studies, fluorescence spectra for **TRapH** (50 μM) were recorded in solvents of different polarities, acetonitrile, methanol, chloroform, dimethylsulfoxide, and aqueous buffer.

**Determination of $pK_a$ of TRapH:**

$pK_a$ of **TRapH** was determined in two ways:

(1) 5% DMSO, aqueous solutions (HEPES, 20 mM) of different pH (2, 3, 4, 4.5, 5, 5.5, 6, 7.4, 8) were prepared either by adding few drops of HCl or NaOH. Solutions of **TRapH** (50 μM) were prepared in 5% DMSO, aqueous buffer (HEPES, 20 mM) at different pH values and absorption spectra were recorded. $pK_a$ of **TRapH** was determined from the absorbance (at 347 nm) vs pH plot using the following Henderson-Hasselbach equation[4]:

$$pH = pK_a + \ log\ ( I_{max} - I/I - I_{min})$$

where $I_{max}$ was obtained from the maximum absorbance value at 347 nm, from the pH dependent absorption titration. $I_{min}$ was obtained from the minimum absorbance value at 347 nm, from the same experiment. I is the absorbance value at 347 nm at a particular pH. $pK_a$ of **TRapH** was obtained from the intercept of the linear fit of the pH vs $\log((I_{max}-I)/(I-I_{min}))$ plot.

(2) Ratios of fluorescence response of **TRapH** at $\lambda_{ex}$ = 347 nm to the fluorescence response of **TRapH** at pH 8 were plotted against pH values. The pH value corresponding to the mid-point of the sigmoidal fit afforded the $pK_a$ for **TRapH**.

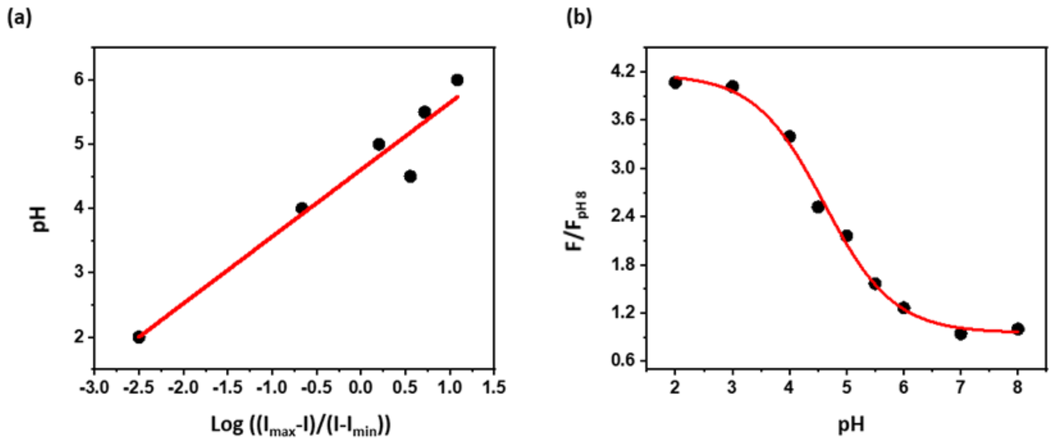


**Figure S6**. (a) pH vs *log* $((I_{max}-I)/(I-I_{min}))$ for **TRapH** was plotted to obtain the $pK_a$ of **TRapH**. (b) Determination of $pK_a$ of **TRapH** from fluorescence response at 468 nm emission wavelength.

**<u>Reversibility of the pH-dependent response of TRapH via absorption measurements:</u>**

Reversibility of the pH dependent response of **TRapH** was verified by shuttling the pH of a solution of **TRapH** (50 μM) in 5% DMSO, aqueous buffer (HEPES, 20 mM) between pH 7.4 and pH 4 by addition of either few drops of HCl or NaOH. A ratio of 438 nm (CT band) and 347 nm (dye to dye band) was taken for each cycle up to the $3^{rd}$ cycle. Experiments were performed in triplicate.

**<u>Reversibility of the pH-dependent response of TRapH via fluorescence spectroscopy:</u>**

Reversibility of the pH dependent fluorescence change of **TRapH** was verified by shuttling the pH of a solution of **TRapH** (50 μM) in 5% DMSO, aqueous buffer (HEPES, 20 mM) between pH 7.4 and 4 either by adding few drops of HCl or NaOH. Fluorescence spectra of **TRapH** (50 μM) were recorded by exciting at both 347 nm and 450 nm. Ratios of emission corresponding to excitation at 450 nm ($\lambda_{em}$ = 538 nm) to that corresponding to excitation at 347 nm ($\lambda_{em}$= 468 nm) were plotted for each cycle till $3^{rd}$ cycle to verify the reversibility of the pH dependent fluorescence response of **TRapH**.

**<u>pH-dependent $^{1}H$ NMR spectra for TRapH:</u>**

Solutions of **TRapH** were prepared in 1:1 DMSO-$d_6$: $D_2O$ (500 μL) solvent system. pH values for the solutions were adjusted to either 3.5 or 7.5 by adding few drops of HCl and NaOH, respectively. $^{1}H$ NMR spectra of both the solutions were recorded at 298 K on 600 MHz spectrometer.

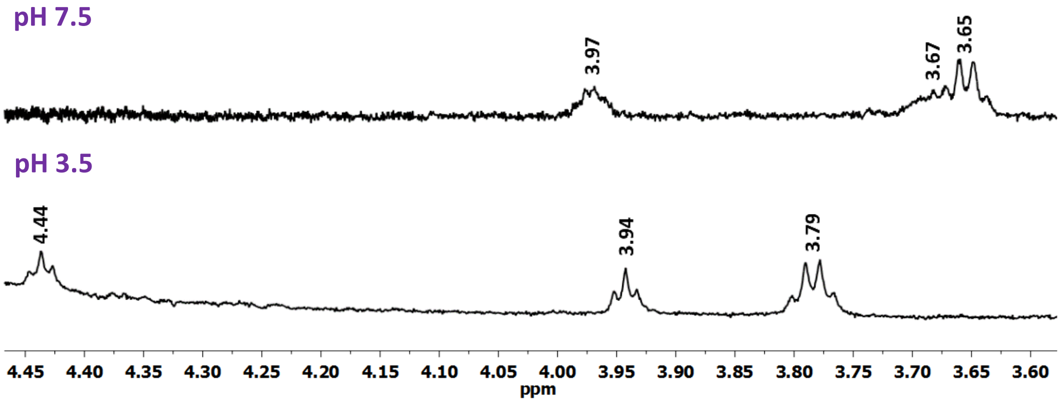


**Figure S7.** Aliphatic region of $^1$H NMR spectra of **TRapH** recorded at two different pH values, pH 7.5 (top panel) and pH 3.5 (bottom panel).

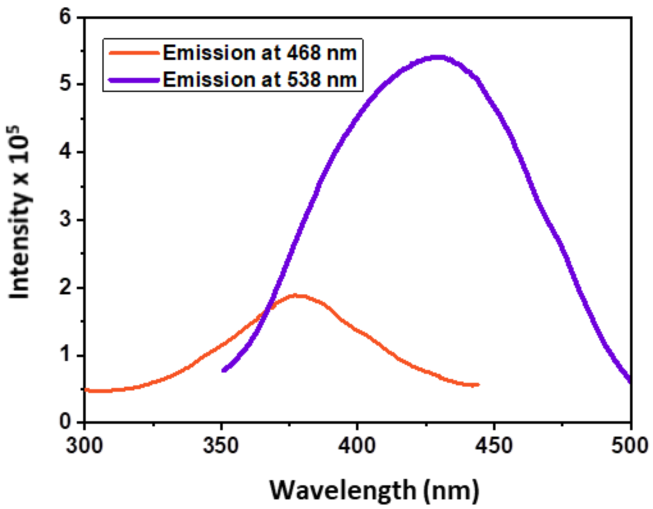


**Figure S8.** Excitation spectra of **TRapH** (50 μM) in 5 % DMSO in aqueous buffer at pH 7.4, corresponding to emission at 468 nm and 538 nm, respectively.

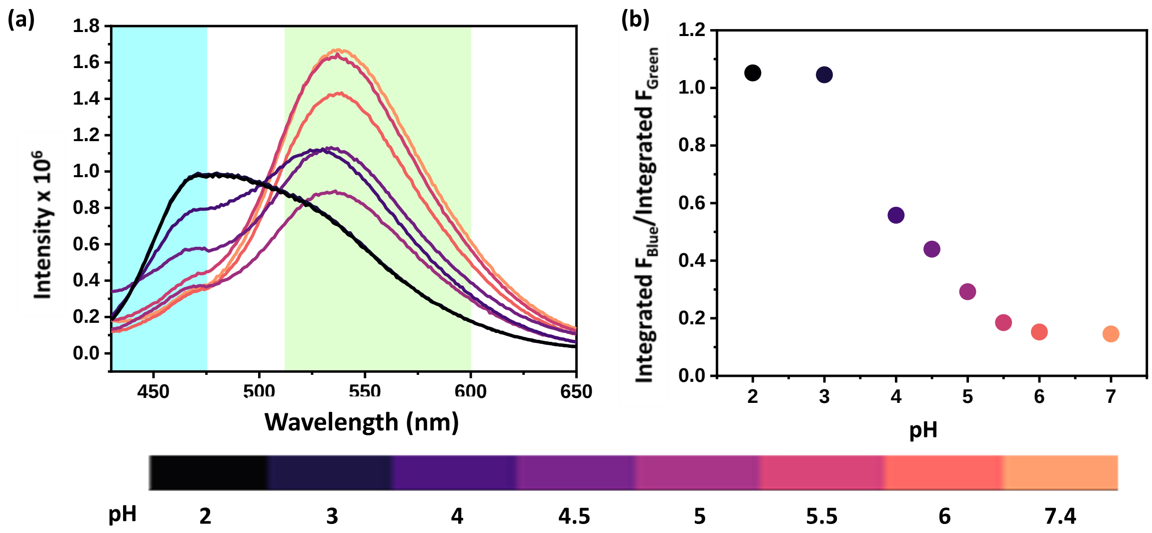


**Figure S9.** (a) Fluorescence response of **TRapH** (50 µM) to pH (2, 3, 4, 4.5, 5, 5.5, 6, 7, 8) in 5% DMSO, aqueous buffer (HEPES, 20 mM), $\lambda_{ex}$ 405 nm (exciting both the bands), slit width 3 nm x 3 nm. (b) Variation of ratios of observed integrated fluorescence intensity ($F_{blue}$) (430-475 nm) over observed integrated fluorescence intensity ($F_{green}$) (510-600 nm) over a pH range of 2 to 8.

### <u>Confocal fluorescence imaging:</u>

HeLa cells were cultured in DMEM (Sigma-Aldrich®), supplemented with Fetal Bovine Serum (10 %, Gibco®), and antibiotic (100x, 10 ml/L) in T25 culture plates at 37 °C under humidified air containing 5 % $CO_2$. For HeLa cells, additional glucose (3.5 mg/ L) was added to the medium. A day before the imaging, the cells were plated on home-made glass coverslip bottomed petriplates (35 mm diameter, Tarsons) coated with polylysine (200 µg/mL). Fluorescence images of the cells were recorded on a confocal microscope (LSM 880, Carl Zeiss, Germany) using 40x oil immersion objectives using 405 nm laser as the excitation source for **TRapH** and emission was collected from 430 to 475 nm for blue channel and from 510 nm to 600 nm for green channel. DMEM media without phenol red (pH adjusted to 7.4 or 4.5) was used during the confocal studies. A stock solution of **TRapH** (10 mM) was prepared in DMSO. The cells were incubated with **TRapH** (10 µM in 5 % DMSO in DMEM media without phenol red pH 7.4) for 5 min at 37 °C under humidified air containing 5 % $CO_2$. After staining, the cells were washed with DMEM media without phenol red pH 7.4 and imaged. To check pH dependent intracellular response of **TRapH**, cells were incubated with **TRapH** (10 µM in 5 % DMSO in DMEM media without phenol red pH 7.4) for 5 min, washed and incubated with **Amphiphilic Peptide 1a**[6] (10 µM in 5 % DMSO in DMEM media without phenol red pH 7.4) for 45 min. After 45 min, cells were washed and an image was recorded. Following this, the media was removed and DMEM media without phenol red pH 4.5 was added. Cells were imaged at intervals of 1 min over 20 min following media change.

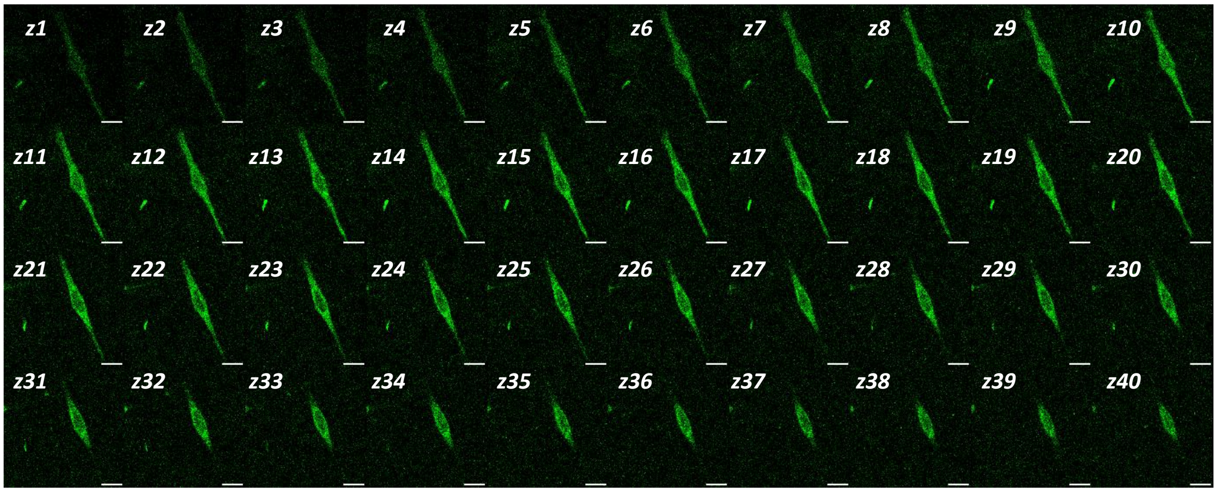


**Figure S10.** Representative confocal single z plane images of same HeLa cell (z1 to z40; 1 µm apart) depicting cell permeability of **TRapH**. Cells were incubated with 10 µM of **TRa-pH** in 5 % DMSO in DMEM media without phenol red (pH 7.4) for 5 min, then washed with DMEM media without phenol red and imaged; $\lambda_{ex}$ = 405 nm; $\lambda_{em}$: 510-600 nm.

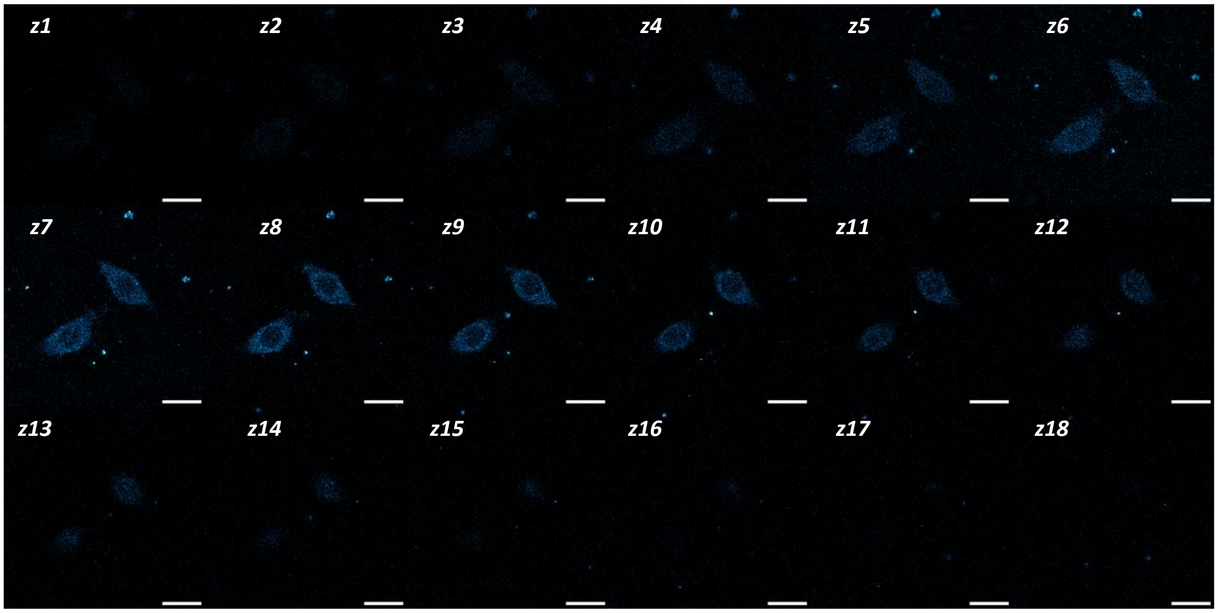


**Figure S11.** Representative confocal single z plane images of same HeLa cells (z1 to z18; 1 µm apart) depicting cell permeability of **TRapH**. Cells were incubated with 10 µM of **TRapH** in 5 % DMSO, DMEM media without phenol red (pH 7.4) for 5 min, then washed with DMEM media without phenol red and imaged; $\lambda_{ex}$ = 405 nm; $\lambda_{em}$: 430-475 nm.

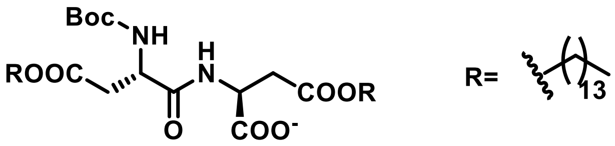


**Figure S12.** Structure of **Amphiphilic peptide 1a.**

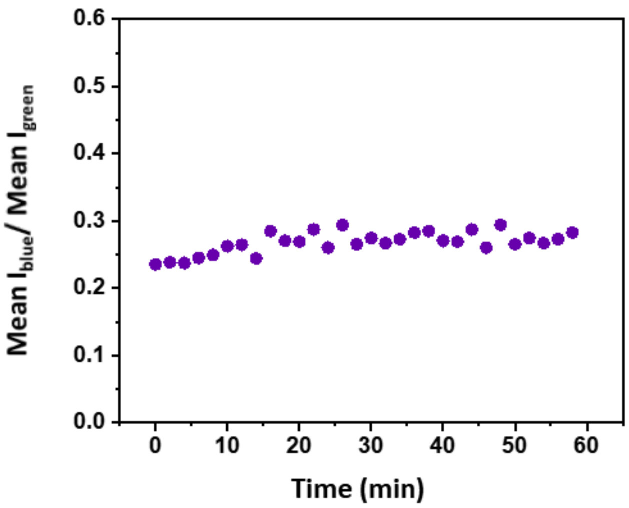


**Figure S13.** Ratio of variation of mean fluorescence intensities of **TRapH** in blue and green channels in HeLa cells with time, depicting photo-stability of **TRapH**. Cells were incubated with 10 µM of **TRapH** in 5 % DMSO in DMEM media without phenol red (pH 7.4) for 5 min, washed with DMEM media without phenol red and then imaged for 1 h at an interval of 2 min. $\lambda_{ex}$ = 405 nm; $\lambda_{em}$ = 430-475 nm for blue channel and 510-600 nm for green channel.

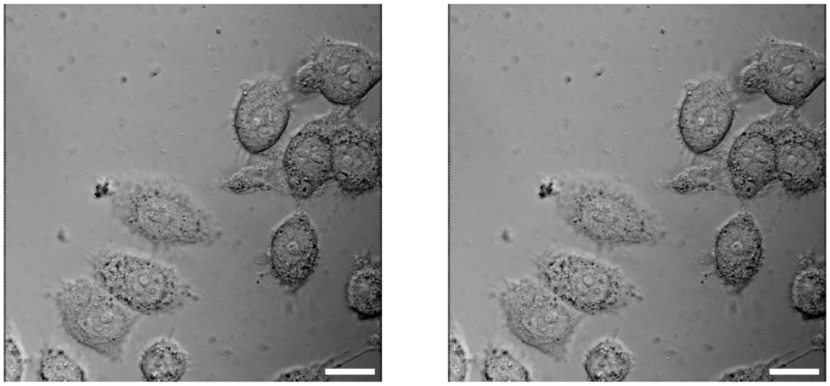


**Figure S14.** Differential contrast images corresponding to Fig. 4a**.**

**Intensity analysis of images:**

For processing and analysis of confocal microscopy images, Fiji (ImageJ, NIH, USA) software was used. For Figure 4b, analysis was done using 10 unique cells from different plates (for each condition). For intensity analysis, regions of interest (ROIs) were drawn around single cells. The intensity of each selection region was obtained using the 'Measure' tool. The intensities in the blue channel in Figure 4a, have been multiplied by 3 for better visualization.

**References:**


1. H. Zong, J. Peng, X.-R. Li, M. Liu, Y. Hu, J. Li, Y. Zang, X. Li and T. D. James, *Chem. Commun.*, 2020, **56**, 515-518.
2. D.-T. Shi, D. Zhou, Y. Zang, J. Li, G.-R. Chen, T. D. James, X.-P. He and H. Tian, *Chem. Commun.*, 2015, **51**, 3653-3655.
3. S. A. Sharber, K.-C. Shih, A. Mann, F. Frausto, T. E. Haas, M.-P. Nieh and S. W. Thomas, *Chem. Sci.*, 2018, **9**, 5415-5426.
4. J. Qian, Y. Xu, X. Qian and S. Zhang, *J. Photochem. Photobiol., A*, 2009, **207**, 181-189.
5. *https://iss.com/resources#fluorescence-quantum-yield-standards*, 2022.
6. S. Kar and N. Madhavan, *Chem. Eur. J.*, 2023, **29**, e202301020.
